# Spatial control of oxygen delivery to 3D cultures alters cancer cell growth and gene expression

**DOI:** 10.1101/522656

**Authors:** William J. Wulftange, Michelle A. Rose, Marcial Garmendia-Cedillos, Davi da Silva, Joanna E. Poprawski, Dhruv Srinivasachar, Taylor Sullivan, Langston Lim, Valery V. Bliskovsky, Matthew D. Hall, Thomas J. Pohida, Robert W. Robey, Nicole Y. Morgan, Michael M. Gottesman

**Affiliations:** Trans-NIH Shared Resources on Biomedical Engineering and Physical Sciences (BEPS), National Institutes of Biomedical Imaging and Bioengineering (NIBIB), National Institutes of Health, Bethesda, MD 20814, USA; Laboratory of Cell Biology, Center for Cancer Research, National Cancer Institute, NIH, Bethesda, MD 20852; Division of Computational Bioscience, Center for Information Technology, NIH, Bethesda, MD 20892; Confocal Microscopy Core Facility, Center for Cancer Research, NCI, NIH, Bethesda, MD 20852; CCR Genomics Core, NCI, NIH, Bethesda, MD 20852; National Center for Advancing Translational Sciences, NIH, Bethesda, MD 20852

**Keywords:** bioreactor, 3D cell culture, oxygen gradient, capillary oxygenation, tumor microenvironment, transcriptome, RNA-seq

## Abstract

Commonly used monolayer cancer cell cultures fail to provide a physiologically relevant environment in terms of oxygen delivery. Here, we describe a three-dimensional bioreactor system where cancer cells are grown in Matrigel in modified 6-well plates. Oxygen is delivered to the cultures through a polydimethylsiloxane (PDMS) membrane at the bottom of the wells, with microfabricated PDMS pillars to control oxygen delivery. The plates receive 3% oxygen from below and 0% oxygen at the top surface of the media, providing a gradient of 3% to 0% oxygen. We compared growth and transcriptional profiles for cancer cells grown in Matrigel in the bioreactor, 3D cultures grown in 21% oxygen, and cells grown in a standard hypoxia chamber at 3% oxygen. Additionally, we compared gene expression of conventional 2D monolayer culture and 3D Matrigel culture in 21% oxygen. We conclude that controlled oxygen delivery may provide a more physiologically relevant 3D system.

## 1. INTRODUCTION

Traditional two-dimensional (2D) cell culture has led to remarkable advances in the study of cancer. However, cancer cells grown in monolayer culture differ significantly from cancer observed *in vivo*, with negative impacts including the predictive value of *in vitro* drug testing. Bridging this gap by better recreating the tumor microenvironment *in vitro* through three-dimensional (3D) cancer models has been an active area of research over the past decade. Cancer cells grown as 3D spheroids compared to 2D monolayer culture have shown increased resistance to chemotherapeutics (Chambers et al., 2014), changes in metabolic signaling pathways (Riedl et al., 2017), and alterations in gene expression (Ghosh et al., 2005).

Common 3D culture models use hydrogels, a class of 3D, bio-compatible polymeric networks, sometimes supplemented with growth factors and other cell nutrients, to support growth of clusters and spheroids of cells in standard multi-well plates. There is general agreement that standard 3D cancer culture models do better at replicating *in vivo* cancer because they maintain important cell-cell and cell-extracellular matrix (ECM) relationships (Haycock, 2011; Pampaloni et al., 2007). This tissue architecture dictates the production and metabolism of growth factors, cytokines, chemokines, and ECM-related molecules (Bissell et al., 2002; Daniel and Smith, 1999) – all important aspects of the tumor microenvironment. Nonetheless, the simple 3D spheroid models fail to accurately reflect normal physiological conditions such as tissue heterogeneity, inter-organ signaling, hypoxic oxygen gradients and nutrient delivery. Larger spheroids frequently develop an hypoxic core due to a lack of vasculature (Edmondson et al., 2014). More complex 3D models employ microfluidics, microfabrication, and multiple cell types in order to address these complexities (Uzel et al., 2016; Vishnubhakthula et al., 2017; Xie et al., 2016). The hope is that complex biological processes such as angiogenesis, cell adhesion, and apoptosis can be more accurately studied through the use of these more advanced 3D culture models, potentially offering clinically relevant insights.

Growing cells in atmospheric oxygen is a common feature among almost all of the extant 2D and 3D culture models, particularly those involving cancer cultures. Usual *in vitro* atmospheres consist of air (~21% O_2_ or 159 mm Hg) that is humidified and supplemented with CO_2_. However, median *p*O_2_ for *in vivo* head, neck, cervical, and breast cancers was estimated to be approximately 10 mm Hg (1.3% O_2_) by Vaupel *et al* (Vaupel et al., 2007). Since the oxygen concentration in cancer tissues dictates metabolic activity, drug sensitivity, and cell survival, supplying cultures with physiologically relevant oxygen levels is essential to increase the relevance of *in vitro* models (Vaupel et al., 1989). To correct this discrepancy, hypoxia models have been developed that use controlled-atmosphere chambers or deferoxamine mesylate salt, a hypoxia-mimetic agent (Heddleston et al., 2009; Sanchez-Elsner et al., 2001). Spatial control of oxygen delivery also appears to be important, as cancer has been shown to respond specifically to gradients of oxygen (Pouyssegur et al., 2006). For example, sarcoma tumors have been shown to migrate more quickly in response to hypoxia gradients, suggesting that oxygen is a key physiological factor when tumors are co-opting vasculature to accommodate new cell growth or hypoxic cores (Lewis et al., 2016). In addition to other microenvironmental factors, oxygen tension and hypoxia-related gene signatures have shown potential to be good indicators of both metastasis and prognosis in human cancer cases (El Guerrab et al., 2017). Since gradients of oxygen are critical in cell differentiation and migration, mimicking such gradients *in vitro* could offer clinically-relevant insights into tumor development, metastasis, and resistance to therapy.

In this study, we examined the effects of controlled oxygen delivery in a 3D bioreactor system. This system extends our earlier work on a single-culture bioreactor (Jaeger et al., 2013), and is comprised of modified 6-well plates with polydimethylsiloxane membranes as the bottom surface. Molded micropillars, mimicking vasculature, spaced 200 μm apart, allow for spatially controlled delivery of oxygen to the culture volume. In the bioreactor, these custom plates sit between a 3% oxygen source, which is delivered through the PDMS membrane into the culture volume, and a top chamber which is maintained at 0% oxygen, thus allowing for a 3%-0% oxygen gradient, and effective delivery of physiological levels of oxygen to the 3D culture volume around the pillars (Jaeger et al., 2013). To compare the effects of this mode of oxygen delivery compared to traditional 2D and 3D culture models, the ovarian cancer and breast cancer cell lines, OVCAR-8 and MCF-7, respectively, were cultured in Matrigel in the bioreactor over 7 and 14 days, after which the cultures were imaged using confocal microscopy. Cells were then harvested and gene expression was analyzed by RNA-seq. Cells in the bioreactor were compared to cells grown in traditional 2D cultures in 21% oxygen, cells grown in 3D culture in Matrigel in 21% oxygen, and cells grown in 3D culture in Matrigel in 3% oxygen. The results show dramatic effects of relative hypoxia (3% O_2_ vs. 21% O_2_) and of oxygen gradients (3% O_2_ to 0% O_2_) on cell growth and gene expression patterns. We also present RNA-seq comparisons of cultures grown in traditional monolayer culture and as spheroids in Matrigel; while the results are consistent with the dramatic differences in gene expression reported in the literature, we believe this to be the first genome-wide comparison of gene expression data between these two commonly used culture conditions.

## 2. MATERIALS AND METHODS

### 2.1 Bioreactor fabrication

The two-chamber bioreactor and custom microplates were designed using the commercially available 3D computer-aided design (CAD) software SolidWorks (Dassault Systèmes SolidWorks Corporation, Waltham, MA). The fabrication equipment included an ABS Dimension Elite 3D printer (Stratasys, Inc., Eden Prairie, MN), an MDX-540 benchtop CNC milling machine (Roland DGA Corporation, Irvine, CA), and the PLS6.150D CO2 laser cutter (Universal Laser Systems, Scottsdale, AZ). Cast acrylic sheets (Marga Cipta, New York, NY) were used to manufacture the CNC parts and to laser cut panels. The panels were bonded using the plexiglass acrylic glue Weld-On 4 (Weld-On, Compton, CA).

### 2.2 Two-chamber bioreactor

The modular bioreactor consists of top and bottom chambers (Figure 1A,D) assembled with ¼” thick white acrylic panels. A ½” thick clear acrylic divider panel, mounted with thumbscrews around the outer perimeter, joins the top and bottom chambers. Super-soft silicone rubber (5963K19 McMaster-Carr) was used as gasket material between the chamber and the divider panel. The bottom chamber’s rear panel has a cutout to mount the sensor and control panel of a ProOx C21 pod (Biospherix, Parish NY). The top chamber’s side panels have input and output gas ports for barbed hose fittings. The divider panel has six cutouts to hold the custom microplates, which have custom-made O-rings made from 1/8” EPDM foam cord (8605K41 McMaster-Carr) glued with Loctite 2011 instant adhesive (Henkel AG & Company, Dusseldorf, Germany) to the outer perimeter. Additionally, the top chamber has an access port on the top panel for fiber optic oxygen sensors (FMI-AP-POF-OX-500, Fluorometrics, Tampa FL) which were read with a NeoFox-GT (Ocean Optics, Largo FL). Front doors made with 1/16” thick clear acrylic were mounted to each chamber using plastic hinges. Rectangular magnets with adhesive backing were used to make the door seal magnetic. Significantly, the front door seals were not completely air tight, in order to allow equilibration of the lower chamber atmosphere to the desired oxygen concentration with the ProOx controller. During an experiment, each of the six cutouts was either filled with a custom microplate or a blank insert made from 1/2” thick clear acrylic. Small weights consisting of non-magnetic 316 stainless steel discs were placed on top of the multi-well plates/blanks to ensure a seal between the two bioreactor chambers.

**FIGURE 1.**
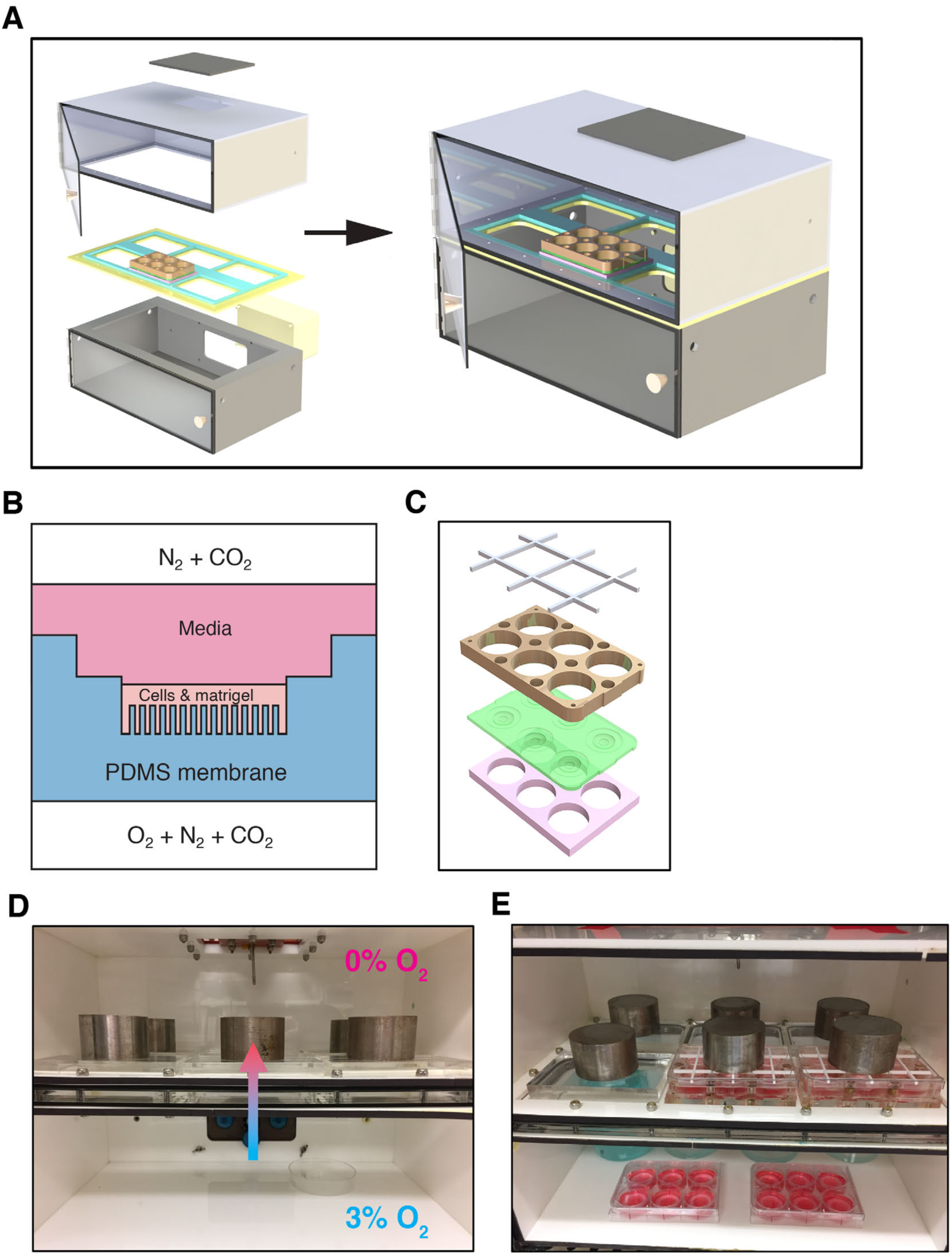
Bioreactor schematics. (a) Exploded view of the dual-chamber bioreactor. (b) Illustration of air-PDMS interface with Cell+Matrigel matrix. (c) Custom PDMS 6-well plates, with magnetic plates shown in blue, PDMS membrane in green, and spacer in white. (d) Diagram of direction of diffusion of oxygen from the 3% O_2_ bottom chamber into the 0% O_2_ top chamber. (e) Photo of bioreactor in use with two gradient plates and two 3% O_2_ plates.

### 2.3 Custom multi-well plates

The culture plates consist of a sandwich, in which the top and bottom pieces are cut with six through-holes and the same footprint as a commercial multi-well plate using a CNC mill, from 3/4” and 3/8” thick clear acrylic blocks respectively. For use, these top and bottom plates are assembled with a PDMS membrane, for which a detailed description follows below. The acrylic plates are held together using 3/8,” and 1/8” diameter N52 axially magnetized disc magnets (K&J Magnetics, Inc., Pipersville, PA) encapsulated in Loctite M31CL epoxy (Henkel AG & Company, Dusseldorf, Germany) for magnetic coupling. The top plate has raised ribs to compress the PDMS membrane around the wells to prevent leaks. The assembled culture plates were used with commercial multi-well plate lids, with a grid-shaped spacer laser cut from 1/8” thick acrylic to allow for faster equilibration with the gas concentrations in the top chamber of the bioreactor. In order to image these custom plates, a 3D-printed insert was also fabricated to fit the custom magnetic plates into the confocal imaging stage. The confocal imaging stage alignment piece was 3D printed in white P430 ABSPLUS plastic (Stratasys, Inc., Eden Prairie, MN).

### 2.4 Membrane fabrication - molds

Negative molds for PDMS membranes were fabricated by combining cast and cut sheets of PDMS with inserts containing micro-wells fabricated using standard soft lithography techniques. Each membrane is the size of a standard multi-well culture plate, with six culture wells per membrane (Figure 1C). All PDMS was fabricated from a Sylgard^®^ 184 Silicone Elastomer kit (Dow Corning Corporation, Midland, MI), except for the PDMS membranes, which also had a layer of Sylgard^®^ 527 applied as later described; base and curing agent were mixed in a 10:1 ratio by weight using a Thinky Mixer ARE-310 (Thinky USA, Laguna Hills, CA), followed by degassing under vacuum and curing on a level plate overnight at room temperature, followed by curing at 80 °C for one hour. Methanol and 2-propanol were purchased from Fisher Scientific (Waltham, MA), and water rinses were done with 18 MOhm water (Millipore, Burlington, MA).

To create the micro-pillar molds, standard SU-8 photolithographic techniques were used. Briefly, a 4” silicon wafer (Nova Electronic Materials, Flower Mound, TX) was baked at 200 °C for 20 min. after which micropillars were patterned using SU-8, a negative epoxy photoresist (MicroChem, Westborough MA). First a thin adhesion layer of SU-8 2002 was spun onto the wafer, prebaked, exposed, and postbaked following manufacturer instructions. Then a layer of SU-8 2150 was applied by spin-coating at 2000 rpm for 25 seconds, baked at 95 °C for 65 minutes, and exposed using a photomask (Jaeger et al., 2013) to pattern micropillars 100 µm in diameter, with 200 µm spacing, over an area of 75 mm x 75 mm. After a postbake with a temperature ramp, and then 25 minutes each at 90 °C and 95 °C, the wafer was agitated gently in SU-8 developer to remove unexposed SU-8, rinsed in isopropanol, water, and dried. Finally, the template was hard baked at 150 °C for 2 minutes, then silanized for 1 h by vapor deposition of (tridecafluoro-1,1,2,2-tetrahydrooctyl)-1-trichlorosilane (UCT Specialties, Bristol, PA) in a vacuum desiccator.

To create a mold for the micro-pillared region of the membranes, a thin 1 mm layer of PDMS was cast on the microfabricated template, then cured at 80 °C, and demolded using methanol to reduce adhesion. 1 cm diameter circles were punched from this patterned sheet using a disposable biopsy punch (Acuderm Inc., Ft. Lauderdale, FL), and measured to ensure each set of six used in a membrane was within 50 µm thickness.

Each full membrane mold was assembled on a borosilicate glass sheet measuring 15.24 x 12.70 x 0.65 cm (McMaster-Carr), with attachment of all the constituent PDMS pieces to the glass and to each other achieved by plasma bonding using a PE-100 Plasma System (Plasma Etch, Inc., Carson City, NV).

A sheet of PDMS measuring 15.24 x 12.70 x 0.36 cm was punched with two rows of 3 circles (Ø = 30 mm), aligned with the positions of the culture wells. A PDMS frame, approximately 8mm wide and aligned with the edges of the glass plate, was added to create a mold depth of approximately 7.0 mm. To create a negative for the mid-well (Figure 1B), 4.5 mm thick cylinders of PDMS (Ø = 18 mm) were bonded to the glass in the center of each of the six holes in the PDMS base layer. The PDMS micropillar molds described above, each 1 cm in diameter and 1.0 mm thick, were attached to the center of these cylinders. Finally, the fully-assembled membrane mold was plasma activated then silanized for 1-3 h by vapor deposition of (tridecafluoro-1,1,2,2-tetrahydrooctyl)-1-trichlorosilane (UCT Specialties, Bristol, PA).

### 2.5 PDMS membrane fabrication

To fabricate PDMS membranes for cell culture, Sylgard^®^ 184 was prepared as earlier described and poured into membrane molds to a depth of 2.5 mm. After curing this layer, the membranes were removed from the mold, and a second, softer layer of PDMS was applied to the membrane surface around the wells. This softer PDMS consisted of a 7:3 (by weight) mixture of Sylgard^®^ 527 Silicone Dielectric Gel (Dow Corning Corporation, Midland, MI) and Sylgard^®^ 184. The two compounds were each prepared separately according to the manufacturer’s instructions, then combined at the desired ratio, mixed and defoamed using the Thinky Mixer ARE-310 (Thinky USA, Laguna Hills, CA). The soft-layer mixture was poured over the membranes, taking care not to cover the well areas, and the membranes were set on a level plate overnight, then cured at 80 °C for 60 minutes. Completed membranes were trimmed, cleaned with isopropanol and water, and then sandwiched between the custom sandwich plates. The assembly was checked for leaks using 2-propanol, then rinsed with water and allowed to dry. Control plates were assembled by excising individual wells from PDMS membranes and attaching them inside standard 6-well plates (Corning Inc., Kennebunk, ME) using 25 μL PDMS. Sets of sandwich and control plates were sealed in plastic and then sterilized with a Cobalt-60 irradiator using 60 krads.

### 2.6 Cell culture and experimental conditions

The OVCAR-8 and MCF-7 cell lines were stably transfected using expression vectors encoding the fluorescent protein *DsRed2* from Clontech (Mountain View, CA) according to previously reported protocols (Brimacombe et al., 2009). Selection of *DsRed2*-expressing cells was enforced with G418 (Corning Life Sciences, Bedford, MA) during passaging and maintenance of cell lines; G418 was not used during bioreactor experiments. Cells were grown in T-75 polystyrene culture flasks for four days prior to the bioreactor experiments, allowing cells to reach approximately 80% confluency. Dulbecco’s Modified Eagle Medium (DMEM) 1X was supplemented with 10% fetal bovine serum (FBS) (Atlanta Biologicals, Lawrenceville, GA), penicillin (100 units/mL), streptomycin (100 µg/mL), and L-glutamine (2 mM), which were purchased from Life Technologies (Carlsbad, CA) and used as received. 0.25% trypsin-EDTA (Gibco, by Life Technologies, Burlington, ON) was used to dissociate cells.

### 2.7 Atmosphere control within the bioreactor

The custom-built bioreactor used in these experiments has two chambers with independent control of gas concentrations. Nitrogen is supplied by filtration of pressurized dry air by an N2-14 Nitrogen Generator (Parker Hannifin Corp., Haverhill, MA); carbon dioxide is supplied centrally by the laboratory building. The top chamber of the bioreactor is maintained at nominally anoxic (< 0.5% O_2_) conditions, with a constant flow of a 95% nitrogen, 5% carbon dioxide mixture through the chamber. The gas concentrations in the bottom chamber are controlled with a ProOx Model C21 (BioSpherix, Parish, NY), and were set at 3% O_2_ and 5% CO_2_ (~92% N_2_) for all the experiments reported here. When first putting the culture plates into the bioreactor, as well as after every media change, both bioreactor chambers were purged with high gas flow until the desired oxygen concentrations were reached, typically for about 15 minutes. The oxygen concentration in the top chamber was monitored with a NeoFox Oxygen Sensing System (Ocean Optics, Dunedin, FL) using a fiber-optic fluorescence lifetime probe. The ProOx controller both measured and controlled the oxygen concentration in the bottom chamber. Both chambers were maintained at the desired gas concentrations for approximately three days prior to the start of experiments.

### 2.8 Cell plating for 3D cultures

Matrigel^®^ Matrix Growth Factor Reduced Basement Membrane (Corning Life Sciences, Bedford, MA) was used to support all the 3D cultures in these experiments. The Matrigel was aliquoted upon receipt, and stored at −20°C until use; aliquots were thawed at 4°C overnight as needed for experiments. Fibronectin (FN) 1 mg/mL solution was purchased from Sigma Aldrich (St. Louis, MO), and diluted to a working concentration of 50 µg/mL in 1X PBS (K-D Medical, Columbia, MD) immediately before use.

To promote adhesion of the Matrigel to the PDMS wells, 150 µL FN solution was added to each well and allowed to remain at room temperature for 60 minutes, after which the excess FN solution was aspirated. Next, the plates were chilled at 4 °C for 15 minutes, and then coated with a thin layer of Matrigel by adding 150 µL of Matrigel diluted in DMEM to a concentration of 0.2 mg/mL in each well, held at room temperature for 60 minutes, and then aspirated off. Plates were then chilled again at 4 °C for a minimum of 15 minutes until cells were ready to be plated. Pipette tips were chilled at –20 °C, and the plates were kept on ice throughout the cell plating steps. An aluminum block with pillars to contact the membrane in each well was used to maintain the wells at 4 °C during Matrigel deposition.

Cells grown on polystyrene were trypsinized and an aliquot was stored in Buffer RLT plus (QIAGEN, Germantown, MD) with 1% β-mercaptoethanol (Sigma Aldrich, St. Louis, MO) for RNA extraction, while the remaining cells were suspended in DMEM and mixed with Matrigel to a concentration of 3 mg mL^-1^ Matrigel and 6 × 10^5^ cells mL^-1^. A total volume of 100 μL of this cell suspension was pipetted into each well. Once plating was completed, plates were put into a 37 °C incubator and flipped every 2.5 mins for 10 mins and then every 5 mins for 35 mins in order to obtain the desired cell distribution while the Matrigel gelled.

After Matrigel gelation, 5 mL of the 10% FBS media, prepared as earlier described, that had been conditioned for approximately 20 hours in 21% O_2_, 3% O_2_, or 0% O_2_, was added to each well on the respective plate. Plated cells were then subjected to one of three conditions. They were cultured in either 1) a standard incubator with 21% O_2_, 2) in the lower chamber of the bioreactor, which contained 3% O_2_, 5% CO_2_, and 92% N_2_ (referred to in this study as 3% O_2_), or 3) at the interface between the upper and lower chambers of the bioreactor, where an O_2_ gradient is created through a membrane (Figure 1D). Media was changed at days 7, 10, and 13, each time with pre-conditioned media. Three open petri dishes with 75 mL of an aqueous CuSO_2_ solution and a beaker with 500 mL of an aqueous CuSO_2_ solution were used to humidify the bottom chamber (Figure 1E). The gas supply for the top chamber was bubbled through water to ensure humidification. The cultured cells were imaged at day 7 or day 14 using confocal microscopy.

### 2.9 Growth curves

The growth of both MCF-7 and OVCAR-8 cell lines was measured for 14 days. 6 x 10^5^ cells per well were plated in both gradient and control 6-well plates. Cells were cultured as previously described, then harvested at days 1, 3, 5, 7, 10, and 14. Plates were removed from the incubator and the media aspirated. Cell Recovery Solution (200 μL, Corning Life Sciences, Bedford, MA) was added to each well and the plates were placed on a rocker at 4 °C for 1 h. The solution from each well was transferred to a separate collection tube, and 1 mL PBS 10X was used to rinse each well to collect the remaining cells. Cells were then counted using a Cellometer™ Auto T4 (Nexelcom Biosciences, Lawrence, MA).

### 2.10 RNA extraction

At the experiment end points of 7 and 14 days, two of the six wells in each plate were excised and cells were extracted from Matrigel by rocking in Corning™ Cell Recovery Solution (Fisher Scientific, Waltham, MA) at 4 °C for 1 h. Wells were rinsed with PBS 1X to collect the remaining cells. The collected cells were then rinsed with 1 mL PBS 1X and centrifuged for 4 mins at 10,000 rpm at 4 °C. Cell pellets were then lysed with RLT Buffer (Qiagen, Hilden, Germany) with 0.01% ß-mercaptoethanol and stored at –80 °C until RNA isolation. Total RNA was isolated from lysed cells according to the protocol supplied with the Qiagen RNeasy^®^ Mini Kit. The RNA integrity and concentration were determined using the Agilent Technologies TapeStation 4200.

### 2.11 Confocal imaging

Confocal imaging of the 3D cultures was done after 7 and 14 days for both cell lines using wells that had not been harvested for RNA. After RNA extraction, the remaining wells were prepared for imaging. Confocal imaging was performed using a Zeiss LSM 710 NLO 2-photon imaging system and Zeiss ZEN software (Carl Zeiss Microscopy, LLC, Thornwood, NJ). For the cells grown in the custom sandwich plates, a 3D-printed adapter was needed to position the plates closer to the microscope objective. PBS 1X with Hoechst^®^ 33342 dye (0.01 mg/mL) (Sigma Aldrich, St. Louis, MO) was used to stain cells prior to image acquisition. Confocal image stacks were rendered in 3D using ImageJ (NIH, Bethesda, MD)(Schneider et al., 2012). To achieve equal aspect ratios for 3D renderings, image sequences were padded with 8-bit grayscale slices with uniform pixel value of 1. Due to differences between gradient and control conditions, different detector gain settings were used for different experiments and digital gain was adjusted accordingly to the darkness of each well. These differing parameters have been accounted for in adjusting the ‘Brightness and Contrast’. The DsRed2 channel brightness and contrast were set to a minimum of “0” and a maximum of “20” for each 14-day image sequence and a maximum of “40” for each 7-day image sequence. The Hoechst 33342 channel brightness and contrast was set to a minimum of “0” and a maximum of “100” for each image sequence.

### 2.12 RNA-seq

Total RNA for 2D, 21%, 3%, and gradient conditions was extracted from the stored samples using a Qiagen RNeasy™ Mini Kit as described above. MCF-7 7-day, 14-day, and OVCAR-8 14- day sequencing libraries were prepared using the Illumina Stranded mRNA kit (Ref: 15056200) and the Illumina NeoPrep™ Library Prep System (Ref: 20003841). OVCAR-8 7-day sequencing libraries were created with a TruSeq Stranded mRNA Library Prep Kit (RS-122-2101). Both cell line sequencing libraries were sequenced using the Illumina NextSeq500. Sequencing reads were trimmed for adapters and low quality bases using Trimmomatic software and the overall quality was verified with FastQC (Andrew, 2010; Bolger et al., 2014). Reads were then aligned with reference human genome hg19 using two-pass STAR alignment, quantified with featureCounts, and normalized with TMM (Dobin et al., 2013; Liao et al., 2014; Robinson and Oshlack, 2010). Differential expression analysis was performed using the EdgeR software package and visualized with ggplot2 (Robinson et al., 2010; Wickham, 2009). Genes were determined to be significantly differentially expressed if there was a log_2_ fold-change ≥ |± 0.6| and a false discovery rate (analogous to an adjusted *p*-value) ≤ 0.05. Gene Ontology analysis of significantly, differentially regulated genes was performed with the GOseq package and redundant Gene Ontology Biological processes were removed using REVIGO with “Small (0.5)” similarity and default parameters (Supek et al., 2011; Young et al., 2010).

## 3. RESULTS

### 3.1 Bioreactor

The bioreactor design presented here (Figure 1A) uses microfabricated polydimethylsiloxane (PDMS) pillars to explore the role of relative hypoxia and oxygen gradients on the growth and gene expression of cultured cancer cells (Figure 1B). The pillared membranes through which 3% oxygen is delivered are designed to recapitulate capillary delivery of oxygen found in tumors in vivo. This system is an expansion of our previously published work, which utilized a single culture well (Jaeger et al., 2013), and which included modelling calculations of the oxygen concentration around the pillars that should also apply to this system. Our two-chamber system allows for higher throughput with the capacity to culture six 6-well plates (Figure 1C) simultaneously under oxygen gradient conditions. Additionally, our system allows for scalability with minimal additional instrumentation by simply increasing the number of wells per plate. The lower chamber of the bioreactor also serves as a standard hypoxia chamber, allowing simultaneous culture of controls in 3% O_2_ (Figure 1D,E).

### 3.2 Cell proliferation

We measured the growth of OVCAR-8 and MCF-7 cells in Matrigel over the course of 14 days in our bioreactor system and compared it to the same lines grown using standard methods for 3D cell culture in Matrigel either kept at 21% oxygen or 3% oxygen (Figure 2). Both cell lines appeared to follow a sigmoidal growth pattern, albeit at different rates, with slower growth observed in the control plates grown under 3% oxygen for seven days or longer. Surprisingly, cells grown under 21% O_2_ and gradient oxygenation in the bioreactor showed very similar growth rates until day 10 of culture. After day 10, OVCAR-8 cells grown at 21% continued to proliferate, whereas growth slowed substantially for the cells grown in the bioreactor (Figure 2A). The MCF-7 line grown at 21% oxygen and in the gradient condition showed similar proliferation for the full experiment duration, for 14 days (Figure 2B). The similarity in growth between 21% O_2_ and gradient conditions, and the dissimilarity between the 3% O_2_ and gradient conditions, seen for both cell lines, was unexpected. Cells grown in the gradient condition received a maximum of 3% O_2_ through the PDMS membrane; the cells in the hypoxic controls experienced the same 3% O_2_ source as a uniform atmosphere, albeit with oxygen delivery top-down through the media. These results indicate that spatial control of oxygen delivery oxygen leads to differences in cancer cell growth compared to cells grown in traditional 3D culture in Matrigel with 3% or 21% oxygen.

**FIGURE 2.**
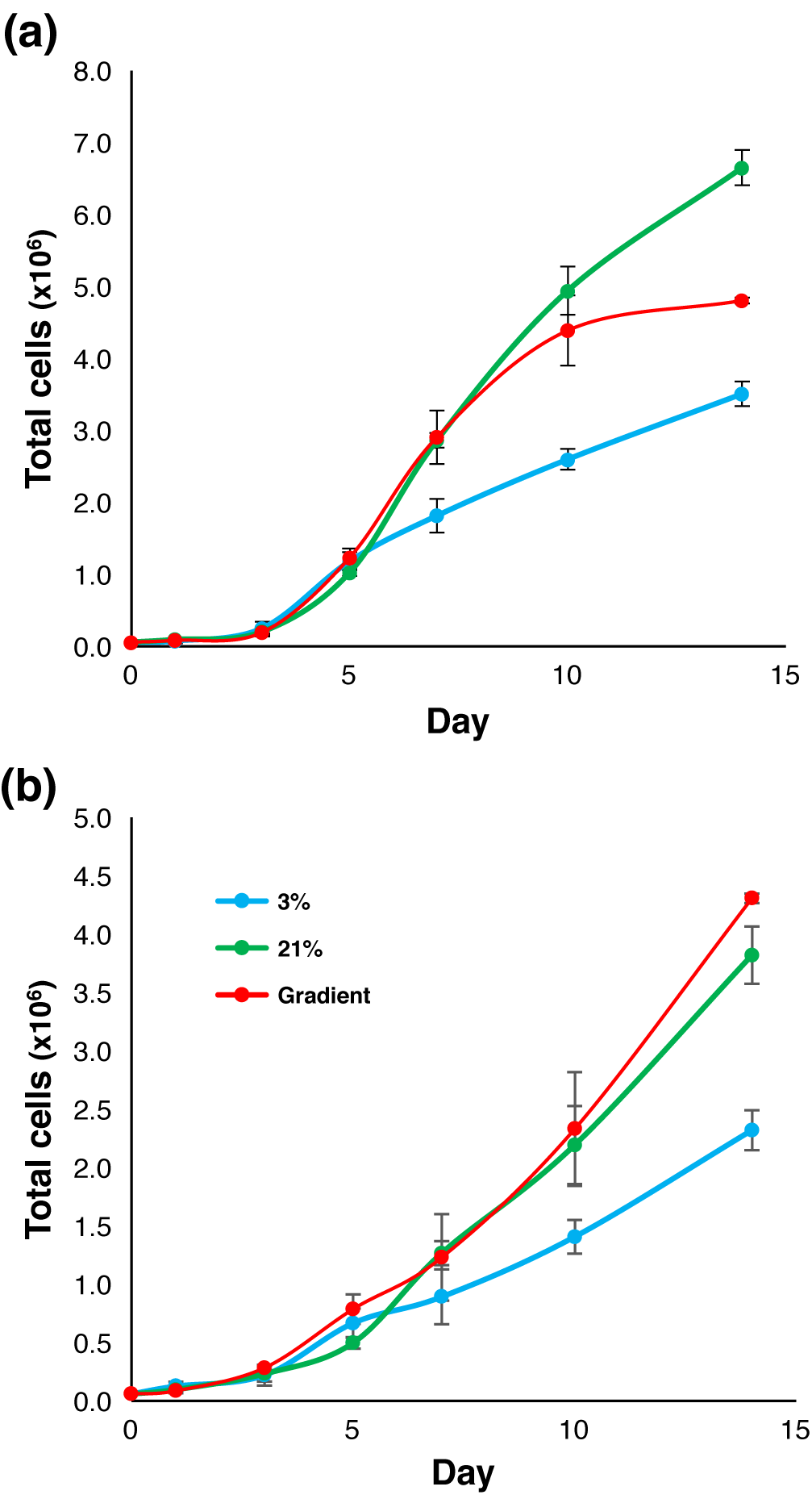
Growth curves of OVCAR-8 and MCF-7 cell lines in 3D Matrigel ECM. (a) OVCAR-8 growth curve illustrating total cell count for cultures grown in various oxygen conditions. (b) MCF-7 growth curve illustrating total cell count for cultures grown in various oxygen conditions.

### 3.3 Imaging of cell growth in the bioreactor

Using a Zeiss 710 NLO (Carl Zeiss Meditec, Dublin, CA), we performed fluorescent confocal microscopy on cell cultures after 7 days of growth and 14 days of growth (Figure 3). Image stacks of cell cultures were taken beginning below the PDMS membrane and extending beyond the top of the Matrigel surface, with slice thicknesses of either 9.4 or 8.9 μm. Z-stack images were then loaded into ImageJ (Schneider et al., 2012) to create three-dimensional renderings, which revealed morphological differences between cells grown under different oxygen conditions. We found that cells cultured in 21% O_2_ tended to grow as spheroids that were randomly dispersed throughout the Matrigel matrix (Figure 3A-D). In contrast, in cultures grown in 3% O_2_ (Figure 3E-H), some spheroids were present, but in much smaller numbers than those observed in 21% O_2_. Additionally, the observed DsRed2 fluorescence in the 3% O_2_ cultures was much less than that seen in the 21% O_2_ cultures, when both were measured relative to the fluorescence of Hoechst 33342 stain (Sigma Aldrich, St. Louis, MO). This suggests that either *DsRed2* translation had been decreased in the hypoxic environment or that cells had died and DNA remained (Supporting Information Figure S1). Cells cultured in the bioreactor (Figure 3I-L) appeared to be much healthier than those grown in 3% O_2_, yet more uniform than those of the 21% O_2_ controls.

**FIGURE 3.**
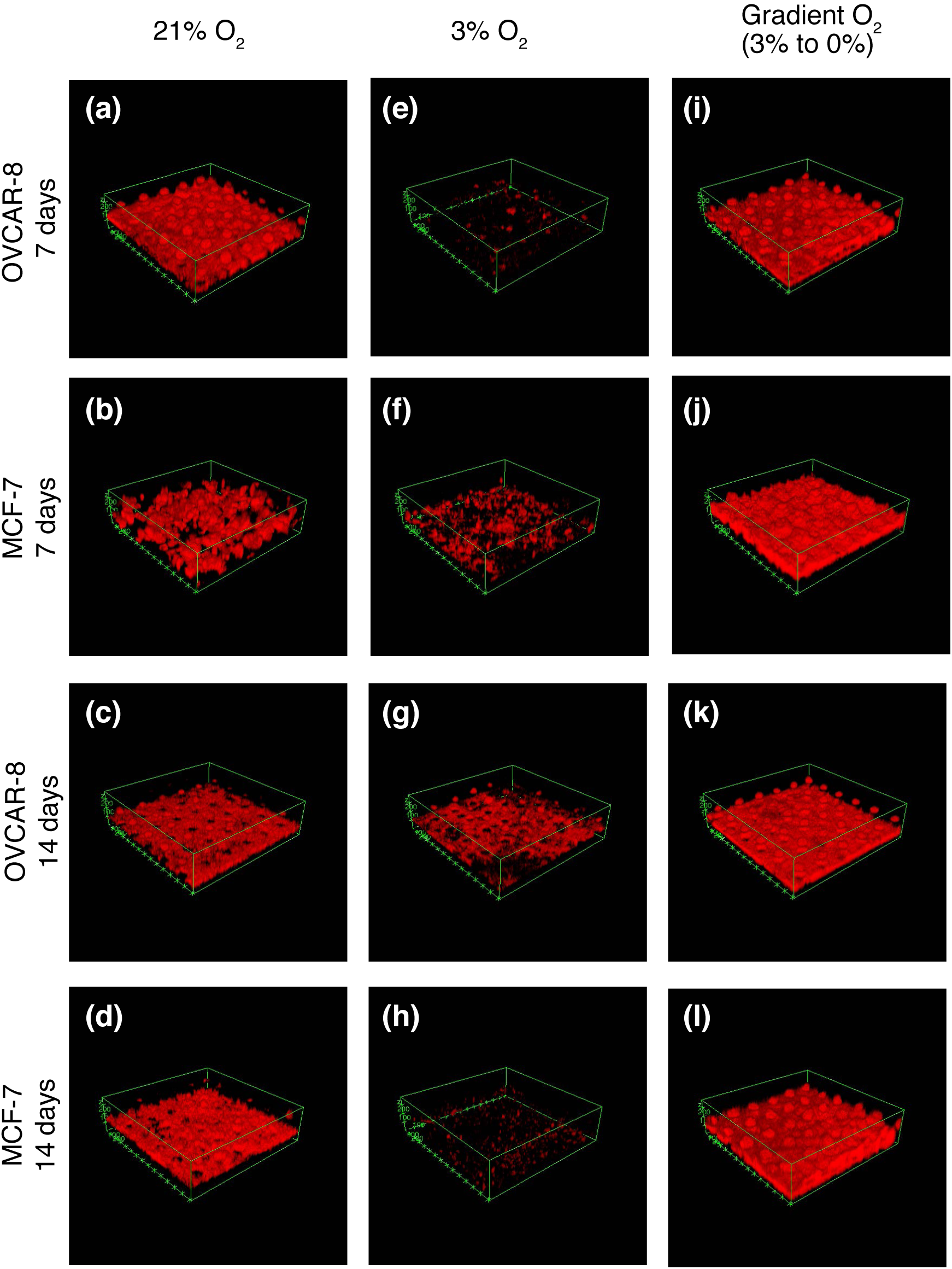
Confocal images of 7-day and 14-day growth of MCF-7 and OVCAR-8 cell lines in 3D Matrigel ECM. (a) OVCAR-8 cells grown in 21% O2 for 7 days. (b) MCF-7 cells grown in 21% O2 for 7 days. (c) OVCAR-8 cells grown in 21% O2 for 14 days. (d) MCF-7 cells grown in 21% O2 for 14 days. (e) OVCAR-8 cells grown in 3% O2 for 7 days, and (f) MCF-7 cells grown in 3% O2 for 7 days. (g) OVCAR-8 cells grown in 3% O2 for 14 days. (h) MCF-7 cells grown in 3% O2 for 14 days. (i) OVCAR-8 cells grown in gradient O2 for 7 days. (j) MCF-7 cells grown in gradient O2 for 7 days. (k) OVCAR-8 cells grown in gradient O2 for 14 days, and (l) MCF-7 cells grown in gradient O2 for 14 days. See also Supplemental Figures 1 and 2.

Cells grown for 14 days in the bioreactor showed slightly different trends in cell morphology. Cells grown in 21% O_2_ exhibited large spheroids extending throughout the Matrigel plugs (Figure 3C,D) for both MCF-7 and OVCAR-8 cell lines. These spheroids were often several hundred microns in diameter, making it difficult to image their entire volumes. The 3% O_2_ controls for both MCF-7 and OVCAR-8 once again showed much less cell growth, but spheroids often extended past the length of the micro-pillars. For both 21% and 3% O_2_ controls, these larger, thicker tissues are not visualized well with *DsRed2* fluorescence, possibly because the large spheroids developed necrotic cores, but light scattering and absorbance at longer working distances and in thick tissues also play a major role in reducing light collection from the top of the spheroids (Supporting Information Figure S2). In contrast, for the 14-day gradient cultures, large spheroids were not nearly as prevalent for either the MCF-7 or OVCAR-8 cell lines. Cells tended to grow in a more uniform layer not more than 100 µm above the pillars from which O_2_ was diffusing (Figure 3K,L).

### 3.4 Analysis of gene expression patterns in cancer cells grown in the bioreactor

#### OVCAR-8

Transcriptome profiles were measured for both cell lines after 7 and 14 days of culture, for the four different culture conditions: 2D monolayer culture with cells in 21% oxygen, 3D cultures grown in Matrigel in uniform 21% O_2_ or 3% O_2_, and 3D cultures grown in Matrigel in the bioreactor gradient condition. Multidimensional scaling of the RNA expression data for all conditions demonstrates clustering of RNA expression patterns (Figure 4) and shows that 2D vs. 3D matrix differences are larger than differences due to oxygenation, as evidenced by sample distances along the abscissa. To discern differences produced solely by 3D vs. 2D culture, we compared cells grown in a standard incubator atmosphere, 21% O_2_ with 5% CO_2_, under two conditions: in 3D in Matrigel and in 2D on tissue culture plastic. We obtained a list of 8,169 differentially expressed genes, 46.96% of all detected genes in our OVCAR-8 7-day dataset (Table 1). These differentially expressed genes were separated by up- and down-regulated genes (Figure 5) and the respectively enriched Gene Ontology (GO) biological processes were investigated (Figure 6). Biological processes involving extracellular matrix organization, ion transmembrane transport, and adhesion were found to be enriched by up-regulated genes while mitotic cell cycle regulation, nitrogen metabolism, and DNA replication were influenced by the down-regulated genes. We found 4,550 differentially expressed genes (27.60% of detected genes) for the OVCAR-8 14-day dataset. Up-regulated genes were involved in ECM organization and inflammatory-related processes. Down-regulated genes were found to be involved in cell division and cell cycle regulation, similar to the 7-day OVCAR-8 21% O_2_ vs. 2D comparison (Figure 6).

**Table 1.** Number of differentially expressed genes (% of total detected)

| **Comparison** | **MCF-7**<br>**7 days** | **MCF-7**<br>**14 days** | **OVCAR-8**<br>**7 days** | **OVCAR-8**<br>**14 days** |
| --- | --- | --- | --- | --- |
| **21% vs. 2D** | 2945 (18.89) | 6780 (43.56) | 8169 (46.96) | 4550 (27.60) |
| **3% vs. 21%** | 828 (5.31) | 1126 (7.23) | 1710 (9.83) | 860 (5.22) |
| **3% vs. gradient** | 470 (3.01) | 1015 (6.52) | 1760 (10.12) | 206 (1.25) |
| **Gradient vs. 21%** | 0 (0.00) | 287 (1.84) | 27 (0.16) | 44 (0.27) |

**FIGURE 4.**
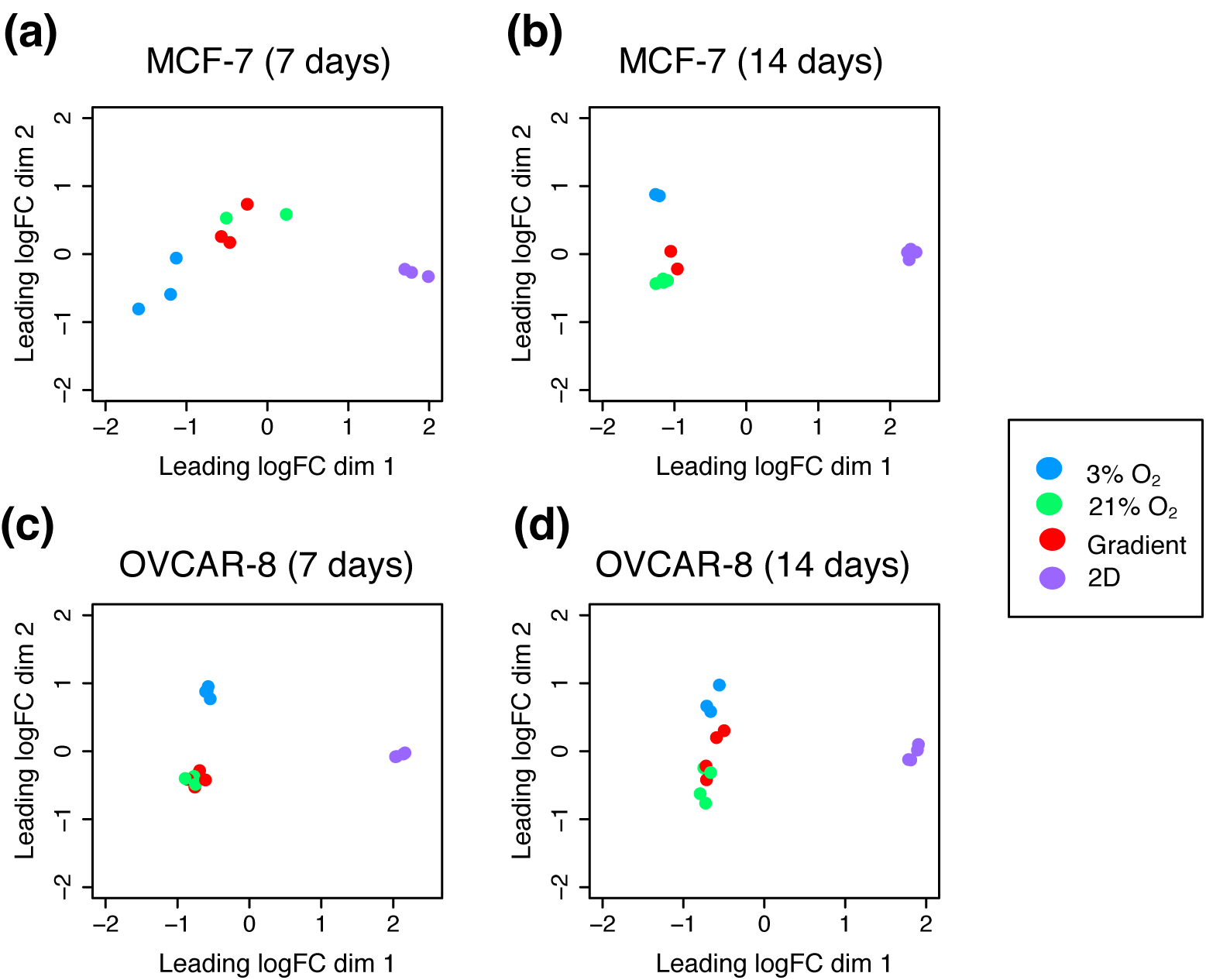
Multi-dimensional scaling (MDS) plots depicting the typical log2 fold changes between genes of (a) MCF-7 7 day dataset, (b) MCF-7 14 day dataset, (c) OVCAR-8 7 day dataset and, (d) OVCAR-8 14 day dataset.

**FIGURE 5.**
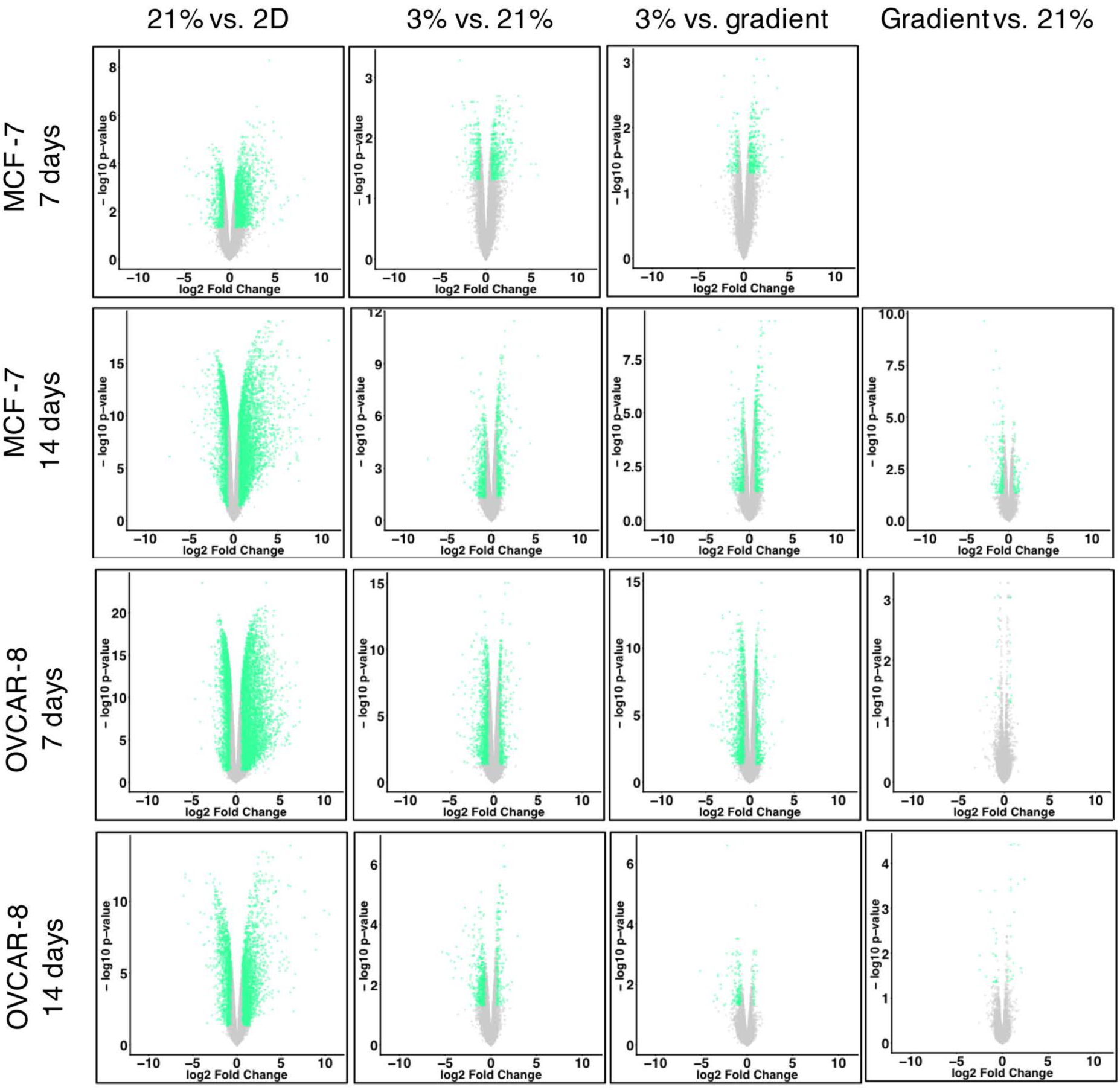
Volcano plots of transcriptomes; genes with a log2 fold change greater than |±0.6| and a False Discovery Rate (FDR) ≤ 0.05 are shown in green, genes not meeting these criteria are grey.

**FIGURE 6.**
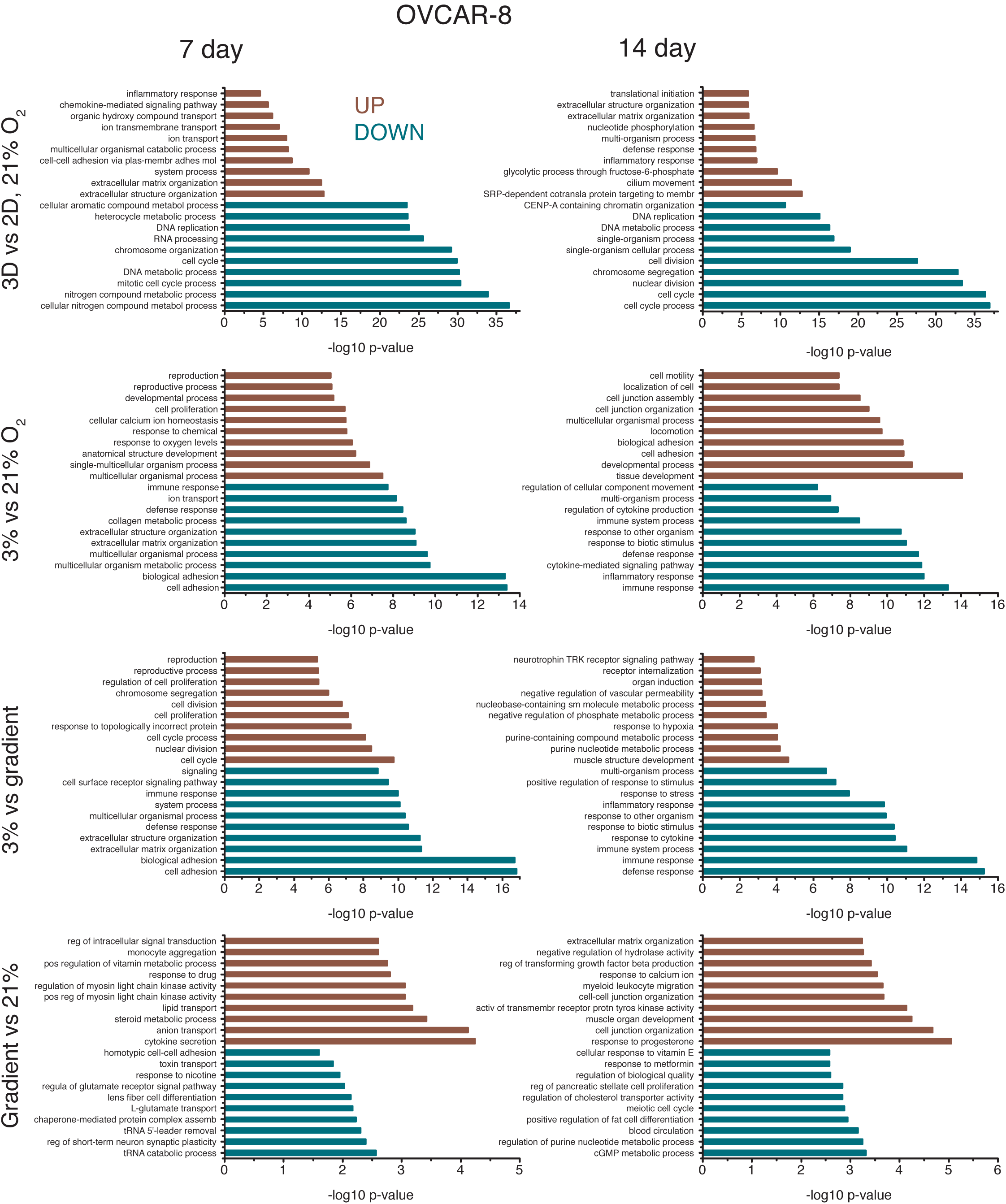
Gene Ontology (GO) biological processes affected by differentially expressed genes for both 7-day and 14-day OVCAR-8 comparisons. Biological processes affected by up-regulated genes are shown in red while processes affected by down-regulated genes are shown in blue. GO biological process lists were generated by GOseq and pruned with REVIGO; the ten processes with the lowest –log10 p-values are shown for 3D vs. 2D 21% O2, 3% vs. 21% O2, 3% vs. gradient, and gradient vs. 21% O2 comparisons.

Our next comparison was of 3% O_2_ cultures to 21% O_2_ cultures to elucidate the effects of hypoxia on 3D Matrigel cultures. Our OVCAR-8 7-day 3% O_2_ vs. 21% O_2_ comparison revealed 1,710 differentially expressed genes (9.83% of total detected) (Table 1). Up-regulated genes were involved in the enrichment of GO processes related to decreased oxygen response and developmental processes, while down-regulated genes were involved in adhesion and ECM organizational processes, all of which are known markers of hypoxia in cancer (Harris, 2002; Wouters and Koritzinsky, 2008). The 14-day comparison revealed that up-regulated genes were involved in the enrichment of tissue and system developmental processes and down-regulated genes were involved in inflammatory and cytokine-related processes (Figure 6).

When comparing our cells grown at 3% O_2_ vs. gradient oxygenation, differences were also observed, despite each condition receiving 3% O_2_, with the gradient condition receiving oxygen through the bottom of the PDMS membrane and the 3% O_2_ control being oxygenated by convection and diffusion through the media covering the cells. OVCAR-8 7-day 3% O_2_ vs. gradient oxygenation resulted in 1,760 differentially expressed genes (10.12% of detected) with up-regulated genes being involved with cell cycle, cell proliferation, and unfolded protein processes. Down-regulated genes for the 7-day comparison were involved with adhesion, ECM organization, and inflammatory response processes. The 3% O_2_ vs. gradient comparison of 14- day data revealed 206 differentially expressed genes (1.25% of detected) with up-regulated genes being involved in several metabolic and hypoxia-related processes and down-regulated genes being involved in cytokine and immune system responses (Figure 6).

Finally, we also compared transcriptomes of our gradient cultures to those of 3D 21% O_2_ cultures. Gradient and 3D 21% O_2_ cultures were found to be very similar in terms of gene expression. As seen in the MDS plots, our gradient and 3D 21% O_2_ samples cluster together for both MCF-7 and OVCAR-8 cell lines (Figure 4). Examination of individual differentially regulated genes showed a similar trend, with gradient vs. 21% O_2_ conditions having the least differences among all comparisons for both MCF-7 and OVCAR-8 cell lines (Table 1). We found only 27 (0.16% of detected) genes differentially expressed in OVCAR-8 cells between the two conditions after 7 days, with the up-regulated genes being involved in interleukin production and lipid and ion transport processes. The down-regulated genes were involved in RNA catabolism and amino acid transport. Similarly, the 14-day gradient vs. 21% O_2_ comparison produced only 44 differentially expressed genes (0.27% of detected). Up-regulated genes were involved in cell junction development, progesterone response, and several migration processes; down- regulated genes were involved in circulatory system and nucleotide metabolism processes (Figure 6). The much greater similarity between the cultures in 21% O_2_ and those in the bioreactor with a 3% O_2_ source, as compared to those cells grown in a 3% O_2_ atmosphere, points to the importance of spatial control of oxygen delivery.

#### MCF-7

The same comparisons were also performed for our MCF-7 7- and 14-day datasets. Initially, our MCF-7 7-day dataset showed a very low amount of significant differential expression between all 3D conditions. However, we believe this low resolution stems from the spread between biological replicates largely due to the effect of a single outlier replicate. We have performed analyses of our 7-day MCF-7 transcriptome data both with and without this replicate. The following data was generated excluding this outlier replicate; analysis including this replicate can be found in Supporting information Figure S3.

Comparison of MCF-7 cultures grown under 21% O_2_ in Matrigel to standard monolayer MCF-7 cultures grown on polystyrene revealed that 3D and 2D cultures exhibit large differences in transcriptome profiles, consistent with what was observed in the OVCAR-8 comparisons (Figure 5) and the separation observed in the MDS plots (Figure 4). Between the 3D and 2D conditions of the MCF-7 7-day data, there were 2,945 genes differentially expressed (18.89% of total detected genes). Enrichment analysis revealed that genes related to biological processes such as ATP and nucleoside metabolic related genes were up-regulated for 3D MCF-7 7-day cultures and genes related to RNA and ribosomal processes were down-regulated compared to the 2D counterpart (Figure 7). Comparing 3D 21% O_2_ transcriptomes to 2D 21% O_2_ for the 14- day MCF-7 data resulted in 6,780 genes (43.56% of detected) that were differentially expressed. Enriched GO biological processes such as ECM organization, locomotion, and receptor signaling pathways were affected by up-regulated genes. Down-regulated genes affected cell cycle and cell division processes and DNA regulation, similar to other 3D vs. 2D comparisons.

**FIGURE 7.**
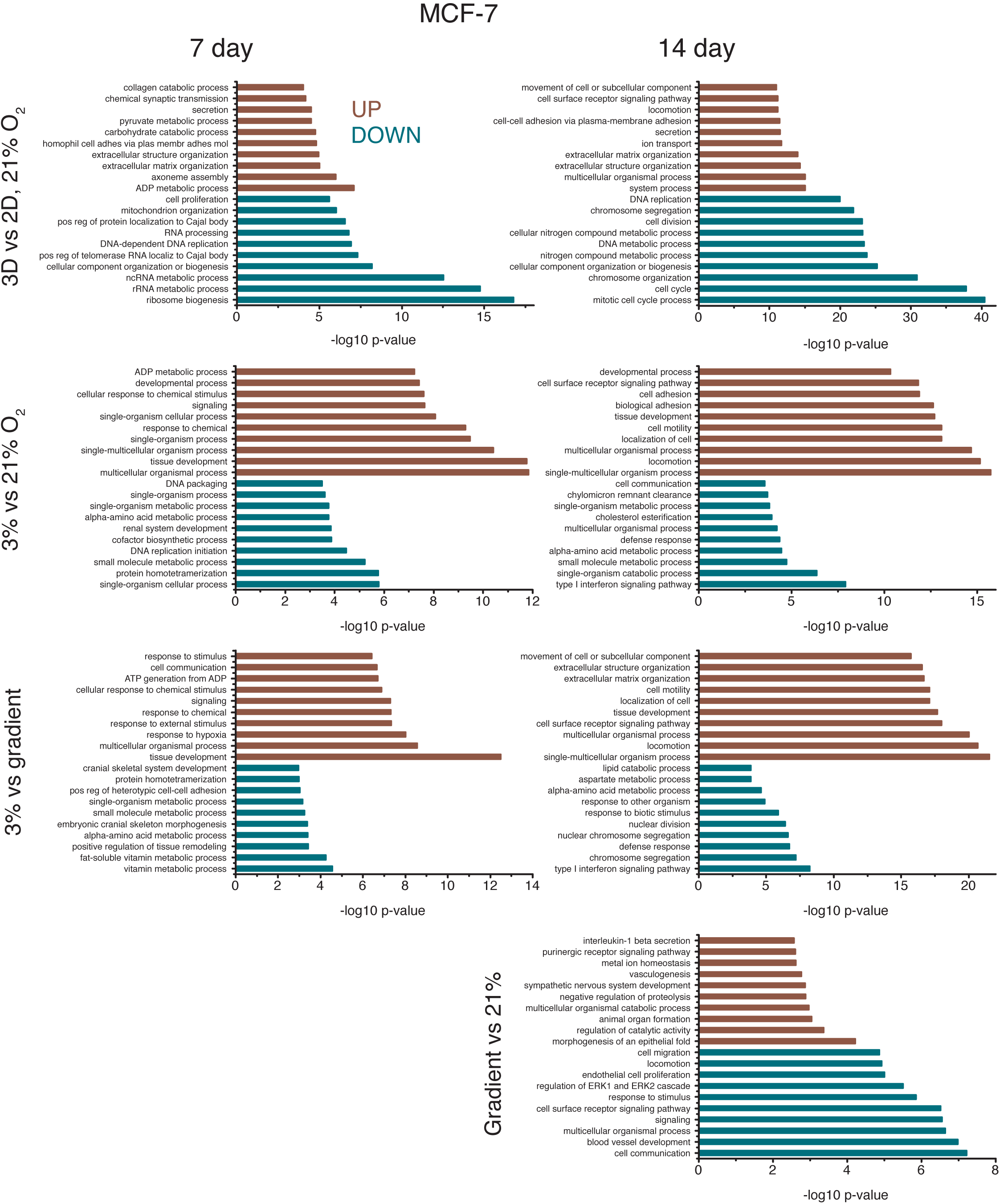
Gene Ontology (GO) biological processes affected by differentially expressed genes for both 7-day and 14-day MCF-7 comparisons. Biological processes affected by up-regulated genes are shown in red while processes affected by down-regulated genes are shown in blue. GO biological process lists were generated by GOseq and pruned with REVIGO; the ten processes with the lowest –log10 p-values are shown for 3D vs. 2D 21%O2, 3% vs. 21% O2, 3% vs. gradient, and gradient vs. 21% O2 comparisons. See also Supplemental Figure 3.

In 3D, as was seen in the OVCAR-8 lines, the comparison of MCF-7 cultures grown under 3% O_2_ vs. 21% O_2_ also showed evidence of hypoxia, with hundreds of differentially expressed genes in the transcriptomes. Examining the MCF-7 7-day data, 828 genes (5.31% of those detected) were differentially expressed between 3% O_2_ and 21% O_2_. Tissue and chemical stimuli as well as organismal processes were enriched by up-regulated genes while biosynthetic and protein-related processes were affected by down-regulated genes. The 14-day MCF-7 3% O_2_ vs. 21% O_2_ comparison revealed localization, adhesion, and motility processes affected by up- regulated genes and type I interferon, metabolism, and cell communication affected by down- regulated genes (Figure 7). Similar to the OVCAR-8 datasets, comparison of 3% O_2_ MCF-7 transcriptomes to the gradient transcriptomes also revealed dissimilarities, reinforcing the idea that the geometric delivery of oxygen is important. Comparison of 3% O_2_ to gradient oxygenation of MCF-7 7-day cultures showed 470 genes (3.01% of detected) differentially expressed, with hypoxia and tissue development processes being enriched by up-regulated genes and metabolic processes being affected by down-regulated genes. The 14-day MCF-7 3% O_2_ vs. gradient comparison produced 1,015 genes (6.52% of detected) being differentially expressed. The up-regulated genes affected migration and motility processes which could be due to cells exposed to 3% O_2_ attempting to move into apical regions of the Matrigel where there is more oxygen. Down-regulated genes affected chromosome segregation and interferon processes (Figure 7).

As was seen in the OVCAR-8 data, comparisons of gradient oxygenation cultures to the 3D 21% O_2_ cultures produced the fewest differences in gene expression for the MCF-7 cells. There were zero differentially expressed genes for the gradient vs. 21% O_2_ 7-day comparison. The same comparison for the 14-day data produced 287 genes (1.84% of detected) that were differentially expressed (Table 1). Several morphogenesis-related GO processes were enriched by the up-regulated genes and blood vessel development and migration processes were affected by the down-regulated genes (Figure 7). Similar to the OVCAR-8 14-day gradient vs. 21% O_2_ comparison, angiogenesis and vasculature processes seem to be affected by genes up- regulated in 3D 21% O_2_ cultures, suggesting there may actually be more hypoxic cells in the cultures grown under ambient atmosphere.

## 4. DISCUSSION

Establishing conditions for the *in vitro* growth of normal tissues and cancer cells that closely mimic physiologic environments offers the promise of growing normal tissues *ex vivo* and of studying the behavior of cancers with the goal of improving therapy. In this work we show that in addition to the profound effect on gene expression of the 3D growth of cancer cells, the mode of oxygen delivery to *in vitro* cultures is an important determinant of cell behavior and gene expression.

There are many ongoing efforts to establish standards for the predictive capacities of established and emerging 3D model technologies, such as lab-on-a-chip devices, in relation to drug-screening and organ mimicry. If *in vitro* models were able to produce results that resemble those of pharmacological studies investigating how drugs are absorbed, distributed, metabolized, and excreted, as well as their toxicity (ADME/TOX), their transition into mainstream usage would likely be expedited. Devices that focus on single-organ models such as the lung-on-a-chip examine cell-cell interactions between two types of cells separated by a porous membrane (Huh et al., 2010). Other models reconstitute multi-organ functions by linking devices together to facilitate metabolic and hormonal signaling between ‘organs’, simulating entire or partial organism physiology (Marx et al., 2016; Pusch et al., 2011; Xiao et al., 2017; Zhang and Radisic, 2017; Zheng et al., 2012). The spatial and temporal delivery of biomolecules in these systems attempts to control the fate of cellular processes with many of these systems utilizing pre-equilibrated media or gas channels to achieve desired oxygenation (Beachley et al., 2015; Grist et al., 2012; Grist et al., 2015; Samorezov and Alsberg, 2015). A standard of systematic oxygenation has not yet been established in these systems, so the implementation of a method of physiological oxygenation to 3D cultures that also allows for high-throughput scalability is therefore highly desirable (Brennan et al., 2014; Place et al., 2017; Sun et al., 2018).

A number of culture models have sought to investigate the effects of oxygen gradients, but scalability into standard cell culture protocols is a pervasive limitation (Adler et al., 2010; Khan et al., 2017; Mehta et al., 2007; Pusch et al., 2011). We report here on a bioreactor system that allows for oxygen delivery through microfabricated PDMS pillars (Figure 1B), providing oxygen delivery to a 3D culture volume at physiological levels (3% O_2_) from pillars spaced similarly to capillaries *in vivo*. We believe this method of oxygen delivery will offer insight into the tumor microenvironment. Our experiments have shown that two cancer cell lines, OVCAR-8 and MCF- 7, thrive when grown in the bioreactor, as evidenced by their growth rate and robust morphology (Figures 2 and 3). Significantly, we did not observe increases in hypoxia-related gene expression even out to 14 days of culture, in contrast to what was seen for the 3D cultures grown at ambient atmosphere (21% O_2_), in which large spheroids developed by day 14. Large spheroids (greater than 200 µm) typically develop hypoxic cores after about 7 days in culture (Däster et al., 2017). In addition, our bioreactor may allow experiments to be designed to thoroughly investigate oxygen-related biological functions such as angiogenesis, cell motility, and metabolic processes, as demonstrated by the GO analysis (Figures 6 and 7).

Recapitulating cell-cell interactions and ECM architecture are well-established advantages that 3D cell culture offers. These microenvironment factors have been shown to dictate homeostasis regulation, injury-response, and cell-signaling (Bissell et al., 2002; Kleinman et al., 2003). The results of our RNA-seq experiments reinforce this notion, illustrating the massive transcriptional differences between cells grown in traditional 2D monolayer and cells grown in 3D using Matrigel at a concentration of 3 mg ml^-1^ (Figure 4). However, we believe there is still a wide gap between 3D cell culture and *in vivo* microenvironments that can be narrowed through proper oxygenation. Simply inducing hypoxic conditions by reducing the amount of available oxygen to *in vivo* venous levels (~23 mm Hg or 3% O_2_) resulted in poor cell growth and morphology (Figure 3E-H) (Cecil et al., 2012). Yet when the same amount of oxygen was delivered basally through our micro-pillared PDMS membrane, both MCF-7 and OVCAR-8 cell lines flourished (Figure 3I-L).

The relatively minimal transcriptional differences between our gradient cultures and the 3D 21% O_2_ cultures was quite surprising, as these conditions differ markedly in maximum oxygen concentration and method of delivery. One explanation for these results is that the proximity of the PDMS micropillars to the cancer cells ensures that there is adequate oxygen delivery to the cells even at 3% oxygen concentrations, whereas the spheroids cultured in an atmosphere with 21% oxygen are no better oxygenated because of the variable depth of the spheroids in the Matrigel, and the limited transport of oxygen to the interiors of the spheroids. Significantly, the cultures grown in 21% oxygen for 14 days exhibit large spheroids that are only weakly fluorescent; it is not known to what extent the cells in these spheroids contribute to the transcriptome data. Another possibility is that the gradient cultures are adapting to low oxygenation, but this adaptation is not revealed through RNA-seq at the 7- or 14-day timepoints. The adaptation may occur during the first few days or even hours of culture and transcriptional equilibrium is reached by day 7. As indicated by our growth curves (Figure 2), the total cell counts of gradient and 3D 21% O_2_ cultures are very similar at day 7, yet start to diverge at approximately day 10; this was observed for both cell lines. Transcriptome analysis of cultures at the 14-day mark revealed more differences between the gradient and 3D 21% O_2_ cultures, but trends of similarity remained the same. We believe our bioreactor provides a novel way of simulating *in vivo* tumor oxygenation by providing spatial control of oxygen delivery to 3D cell cultures in a simply scalable way.

This bioreactor system is an expansion of previous work in which we presented a single-chamber version of the system. The current design allows for six 6-well plates to be cultured under gradient oxygenation, which is a step towards higher throughput screening using this technology. Additionally, the dual-chambered bioreactor could be further adapted for larger multi-well plates with ease since the measurements of the plate inserts resemble standard polystyrene multi-well plate dimensions. The system is also not limited to the discussed gas compositions – using different gas sources would allow for any number of gaseous mixture gradients to be utilized. The bioreactor’s tunability gives the system potential to be adapted for various cell and tissue types.

In summary, our improved bioreactor system allows for spatial control of oxygen delivery to 3D cell cultures via microfabricated PDMS pillars that that were designed to mimic a capillary bed. Our system allows for six 6-well plates to be cultured simultaneously, which greatly increases the throughput from our previous system. We have analyzed our human epithelial cancer cultures with confocal microscopy and RNA-seq to elucidate the differences between 2D and 3D cell culture, as well as 3D cell cultures with different oxygenation conditions. Our results suggest that both MCF-7 and OVCAR-8 cell lines flourish in terms of proliferation and morphology when oxygenation is delivered basally through microfabricated PDMS pillars. This work suggests that not only the amount of oxygen is relevant in cell culture, but also how it is delivered. This insight is relevant when considering *in vitro* models that are designed to provide physiologically relevant results, such as those used in drug-screening and tissue construction.

## Author contributions

MMG, NYM, TJP, RWR, and MDH conceived of the bioreactor system. WJW, MAR, DdS, JEP, and DS developed experimental protocols. NYM, TJP, and MG-C worked on design of the bioreactor. TS and MG-C assisted with fabrication of bioreactor components. MMG, NYM, TJP, RWR, and WJW designed the experiments. WJW performed culture experiments and RNA isolation. LL assisted with confocal microscopy imaging and analysis. VVB performed RNA-seq measurements. WJW performed the RNA-seq data analysis. WJW, NYM, MMG, interpreted data. WJW, NYM, MMG and RWR contributed to writing the manuscript.

## SUPPORTING INFORMATION

Additional supporting information may be found online, Supplemental Figures 1-3.

## Acknowledgements

This research was supported by the Intramural Research Program of the National Institutes of Health. We thank the CCR Collaborative Bioinformatics Resource for allowing us to use their RNA-seq Pipeliner. We also thank the CCR Genomics Core and CCR Sequencing Facility for their assistance with RNA-seq measurements. We are also grateful to George Leiman in the Laboratory of Cell Biology for editorial assistance.

