## Supplemental Figures 1-3 for "Spatial control of oxygen delivery to 3D cultures alters cancer cell growth and gene expression"

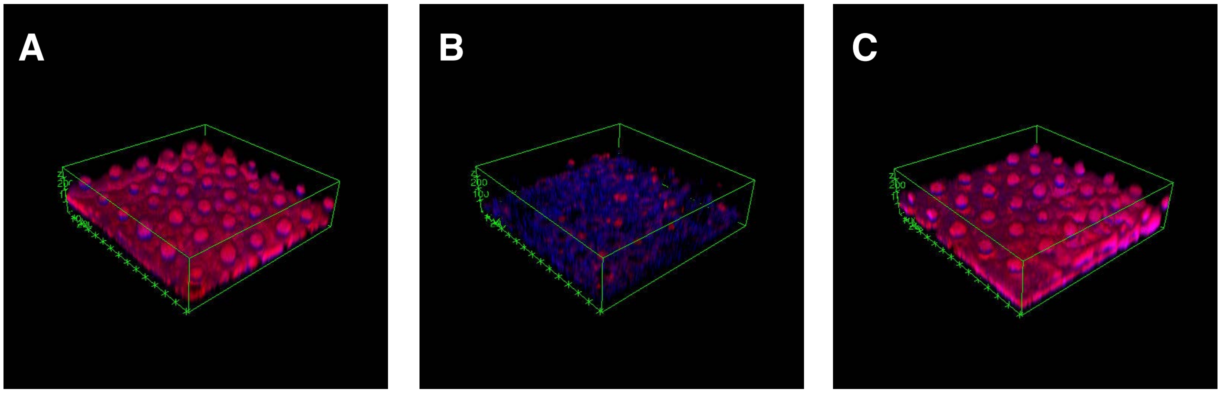


**Supplemental Figure 1:** Confocal images of 7-day growth OVCAR-8 cell lines in 3D Matrigel ECM

Red fluorescence is generated from expression of DsRed2 proteins and blue fluorescence is from Hoechst 33342 staining of DNA. A) OVCAR-8 cells grown in 21% $O_2$, B) in 3% $O_2$, C) and in gradient $O_2$.

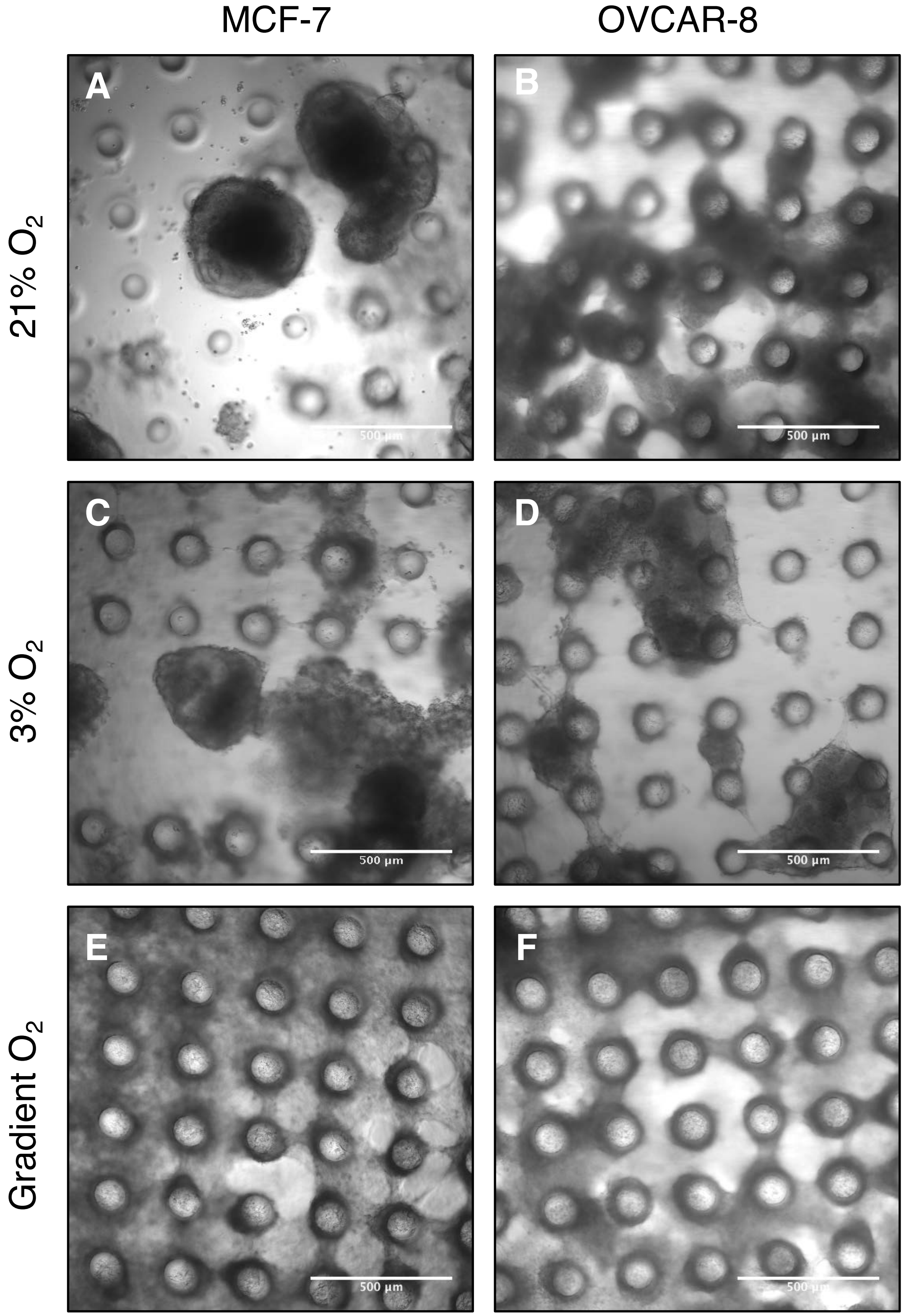


**Supplemental Figure 2:** Brightfield images of 14-day growth conditions

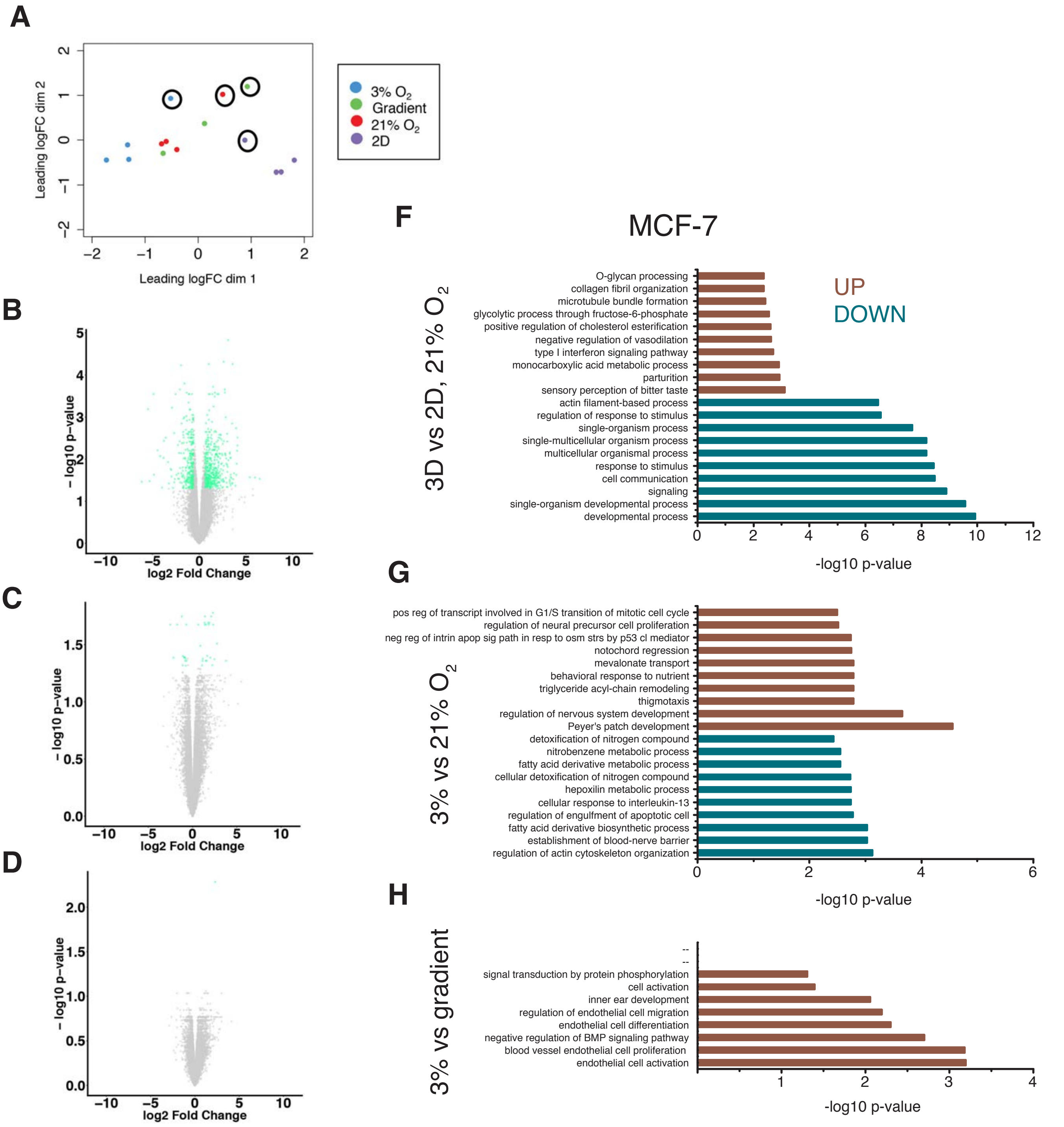


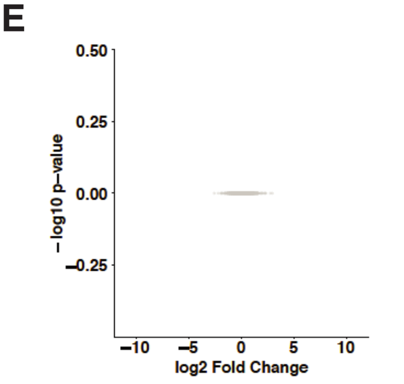


**Supplemental Figure 3:** Visualization of transcriptome comparisons for MCF-7 cell cultures, including outlier replicate

(A) MDS plot depicting the Euclidean distance between log2 fold changes of genes between MCF-7 samples, outlier replicate samples are circled. (B) Volcano plot of transcriptomes for MCF-7 21% $O_2$ vs. 2D comparison; genes with a log2 fold change greater than |±0.6| and a False Discovery Rate (FDR) ≤ 0.05 are shown in green, genes not meeting these criteria are grey. (C) Volcano plots of OVCAR-8 3% vs. 21% $O_2$ , (D) 3% vs. gradient, and (E) gradient vs. 21% $O_2$. (F) Top 10 up-regulated (red) and down-regulated (blue) Gene Ontology (GO) Biological Processes for MCF-7 21% $O_2$ vs. 2D comparison, (G) 3% vs. 21% $O_2$, and (H) 3% $O_2$ vs. gradient.
